# Gene expression inference with deep learning

**DOI:** 10.1101/034421

**Authors:** Yifei Chen, Yi Li, Rajiv Narayan, Aravind Subramanian, Xiaohui Xie

## Abstract

**Motivation:** Large-scale gene expression profiling has been widely used to characterize cellular states in response to various disease conditions, genetic perturbations, etc. Although the cost of whole-genome expression profiles has been dropping steadily, generating a compendium of expression profiling over thousands of samples is still very expensive. Recognizing that gene expressions are often highly correlated, researchers from the NIH LINCS program have developed a cost-effective strategy of profiling only ῀1,000 carefully selected landmark genes and relying on computational methods to infer the expression of remaining target genes. However, the computational approach adopted by the LINCS program is currently based on linear regression, limiting its accuracy since it does not capture complex nonlinear relationship between expression of genes.

**Results:** We present a deep learning method (abbreviated as D-GEX) to infer the expression of target genes from the expression of landmark genes. We used the microarray-based GEO dataset, consisting of 111K expression profiles, to train our model and compare its performance to those from other methods. In terms of mean absolute error averaged across all genes, deep learning significantly outperforms linear regression with 15.33% relative improvement. A gene-wise comparative analysis shows that deep learning achieves lower error than linear regression in 99.97% of the target genes. We also tested the performance of our learned model on an independent RNA-Seq-based GTEx dataset, which consists of 2,921 expression profiles. Deep learning still outperforms linear regression with 6.57% relative improvement, and achieves lower error in 81.31% of the target genes.

**Availability:** D-GEX is available at https://github.com/uci-cbcl/D-GEX.

## 1 Introduction

A fundamental problem in molecular biology is to characterize the gene expression patterns of cells under various biological states. Gene expression profiling has been historically adopted as the tool to capture the gene expression patterns in cellular responses to diseases, genetic perturbations and drug treatments. The Connectivity Map (CMap) project was launched to create a large reference collection of such patterns and has discovered small molecules that are functionally connected using expression pattern-matching (e.g., HDAC inhibitors and estrogen receptor modulators) [1].

Although recent technological advances, whole-genome gene expression profiling is still too expensive to be used by typical academic labs to generate a compendium of gene expression over a large number of conditions, such as large chemical libraries, genome-wide RNAi screening and genetic perturbations. The initial phase of the CMap project produced only 564 genome-wide gene expression profiles using Affymetrix GeneChip microarrays [1].

Despite the large number of genes (῀22,000) across the whole human genome, most of their expression profiles are known to be highly correlated. Systems biologists have leveraged this idea to construct gene regulatory networks and to identify regulator and target genes [2]. Researchers from the LINCS program (http://www.lincsproject.org/) analyzed the gene expression profiles from the CMap data using principal component analysis. They found that a set of ῀1,000 carefully chosen genes can capture approximately 80% of the information in the CMap data (http://support.lincscloud.org/hc/en-us/articles/202092616-The-Landmark-Genes). Motivated by this observation, researchers have developed the L1000 Luminex bead technology to measure the expression profiles of these ῀1,000 genes, called the *landmark genes* (http://support.lincscloud.org/hc/en-us/articles/202092616-The-Landmark-Genes), with a much lower cost (῀$5 per profile) [3]. Therefore, researchers can use the expression signatures of landmark genes to characterize the cellular states of samples under various experimental conditions. If researchers are interested in the expression of a specific gene other than landmark genes, the expression profiles of the remaining ῀21,000 genes, called the *target genes*, can be then computationally inferred based on landmark genes and existing expression profiles. With the L1000 technology, the LINCS program has generated ῀1.3 million gene expression profiles under a variety of experimental conditions.

However, computationally inferring the expression profiles of target genes based on landmark genes is challenging. It is essentially a large scale multi-task machine learning problem, with the target dimension (῀21,000) significantly greater than the feature dimension (῀1,000). The LINCS program currently adopts linear regression as the inference method, which trains regression models independently for each target gene based on the Gene Expression Omnibus (GEO) [4] data. While linear regression is highly scalable, it inevitably ignores the nonlinearity within gene expression profiles that has been observed [5]. Kernel machines can represent dexterous nonlinear patterns and have been applied to similar problems [6]. Unfortunately, they suffer from poor scalability to growing data size. Thus, a machine learning method enjoying both scalability and rich representability is ideal for large scale multi-task gene expression inference.

Recent successes in deep learning on many machine learning tasks have demonstrated its power in learning hierarchical nonlinear patterns on large scale datasets [7]. Deep learning in general refers to methods that learn a hierarchical representation of the data through multiple layers of abstraction (e.g. multi-layer feedforward neural networks). A number of new techniques have been developed recently in deep learning, including the deployment of General-Purpose Computing on Graphics Processing Units (GPGPU) [8, 9], new training methodologies, such as dropout training [10, 11] and momentum method [12]. With these advances, deep learning has achieved state-of-the-art performances on a wide range of applications, both in traditional machine learning tasks such as computer vision [13], natural language processing [14], speech recognition [15], and in natural science applications such as exotic particles detection [16], protein structure prediction [17], RNA splicing prediction [18] and pathogenic variants identification [19].

Here we present a deep learning method for gene expression inference (D-GEX). D-GEX is a multi-task multi-layer feedforward neural network. We evaluated the performances of D-GEX, linear regression (with and without different regularizations) and k-nearest neighbor (KNN) regression on two types of expression data, the microarray expression data from the GEO and the RNA-Seq expression data from the Genotype-Tissue Expression (GTEx) project [20, 21]. GPU computing was used to accelerate neural network training so that we were able to evaluate a series of neural networks with different architectures. Results on the GEO data show that D-GEX consistently outperforms other methods in terms of prediction accuracy. Results on the GTEx data further demonstrate D-GEX, combined with the dropout regularization technique, achieves the best performance even where training and prediction were performed on datasets obtained from different platforms (microarray verse RNA-Seq). Such cross platforms generalizability implies the great potential of D-GEX to be applied to the LINCS program where training and prediction were also done separately on the microarray data and the L1000 data. Finally, we attempted to explore the internal structures of the learned neural networks with two different strategies and tried to interpret the advantages of deep learning compared to linear regression.

## 2 Methods

In this section, we first introduce three expression datasets we used in this study and formulate gene expression inference as a supervised learning problem. We then present D-GEX for this problem and explain a few key deep learning techniques to train D-GEX. Finally, we introduce several common machine learning methods that we used to compare with D-GEX.

### 2.1 Datasets

1. *The GEO expression data* was curated by the Broad Institute from the publicly available GEO database. It consists of 129,158 gene expression profiles from the Affymetrix microarray platform. Each profile comprises of 22,268 probes, corresponding to the 978 landmark genes and the 21,290 target genes. The original GEO data was accessed from the LINCS Cloud (http://www.lincscloud.org/), which has been quantile normalized into a numerical range between 4 and 15. Some of the expression profiles in the GEO dataset are biological or technical replicates. To avoid complications in the learning procedure, we removed duplicated samples (see Supplementary), leaving 111,009 profiles in the end.

2. *The GTEx expression data* consists of 2,921 gene expression profiles of various tissue samples obtained from the Illumina RNA-Seq platform [21]. The expression level of each gene was measured based on Gencode V12 annotations [21] in the format of Reads Per Kilobase per Million (RPKM).

3. *The 1000 Genomes expression data* consists of 462 gene expression profiles of lymphoblastoid cell line samples from the Illumina RNA-Seq platform [22]. The expression level of each gene was also measured based on Gencode V12 annotations [22] in the format of RPKM.

Since the gene expression values of the microarray platform and the RNA-Seq platform were measured in different units (probes vs Gencode annotations) and different numerical scales, we quantile normalized the three expression datasets jointly to retain the maximum information cross platforms. Because one Gencode annotation may include multiple microarray probes, 943 landmark genes and 9,520 target genes in terms of Gencode annotations were left after joint quantile normalization. Details of joint quantile normalization are given in Supplementary. Finally, all the datasets were standardized by subtracting the mean and dividing by the standard deviation of each gene.

### 2.2 Gene expression inference as multi-task regression

Assume there are *L* landmark genes, *T* target genes, and *N* training samples (i.e. profiles); the training dataset is expressed as 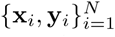, where 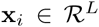 denotes the expression values of landmark genes and 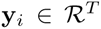 denotes the expression values of target genes in the *i*-th sample. Our goal is to infer the functional mapping 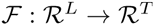 that fits 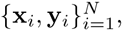, which can be viewed as a multi-task regression problem.

We use Mean Absolute Error (MAE) to evaluate the predictive performance at each target gene *t*,

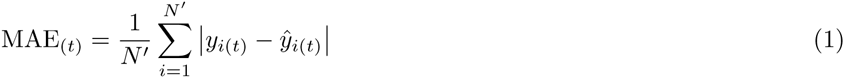

where *N*′ is the number of testing samples and 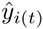 is the predicted expression value for target gene *t* in sample *i*. We define the overall error as the average MAE over all target genes, and use it to evaluate the general predictive performance.

For the microarray platform, we used the GEO data for training, validation and testing. Specifically, we randomly partitioned the GEO data into ῀80% for training (88,807 samples denoted as GEO-tr), ῀10% for validation (11,101 samples denoted as GEO-va) and ῀10% for testing (11,101 samples denoted as GEO-te). The validation data GEO-va was used to do model selection and parameter tuning for all the methods.

For the RNA-Seq platform, we used GEO-tr for training, the 1000 Genomes data for validation (denoted as 1000G-va), and the GTEx data for testing (denoted as GTEx-te). The validation data 1000G-va was used to do model selection and parameter tuning for all the methods.

### 2.3 D-GEX

D-GEX is a multi-task multi-layer feedforward neural network. It consists of one input layer, one or multiple hidden layers, and one output layer. All the hidden layers have the same number of hidden unites. Units between layers are all fully connected. A hidden unit *j* in layer *l* takes the sum of weighted outputs plus the bias from the previous layer *l* − 1 as the input, and produces a single output 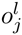 using a nonlinear activation function *f*.

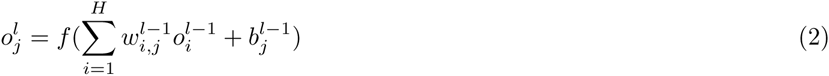

*H* is the number of hidden units. 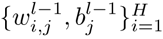 are the weights and the bias associated with unit *j* that need to be learned. We adopt the hyperbolic tangent (TANH) activation function to hidden units, which naturally captures the nonlinear patterns within the data. Linear activation function is applied to output units for the regression purpose. The loss function for training is the sum of mean squared error at each output unit, namely,

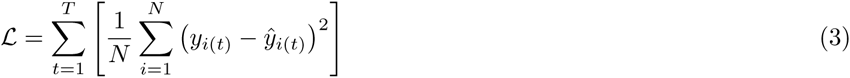

D-GEX contains 943 units in the input layer corresponding to the 943 landmark genes. Ideally, we should also configure D-GEX with 9,520 units in the output layer corresponding to the 9,520 target genes. However, each of our GPUs has only 6 GB of memory, thus we cannot configure hidden layers with sufficient number of hidden units if all the target genes are included in one output layer. Therefore, we randomly partitioned the 9,520 target genes into 2 sets that each contains 4,760 target genes. We then built 2 separate neural networks with each output layer corresponding to one half of the target genes. With this constraint, we were able to build a series of different architectures containing 1῀3 hidden layers each and each hidden layer contains 3,000, 6,000 or 9,000 hidden units. Supplementary Figure S1 shows an example architecture of D-GEX with 3 hidden layers.

Training D-GEX follows the standard back-propagation algorithm [23] and mini-batch gradient descent, supplemented with advanced deep learning techniques. Detailed parameter configurations are given in Supplementary Table S1. For more descriptions about neural networks and their background please see [24]. We discuss a few key training techniques as follows:

1. *Dropout* is a technique to perform model averaging and regularization [10] for neural networks. At the training time, each unit along with its edges is temporarily dropped out with probability *p* for each training sample. Then the forward- and back-propagation are performed on a particularly “thinned” network. For an architecture with *n* units performing dropout, there are 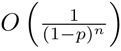 such thinned networks. At the testing time, all the units are retained with weights multiplied by 1 − *p*. Therefore, dropout can be seen as model averaging of exponentially many different neural networks in an approximate but efficient framework. Dropout has been shown to suppress co-adaptation among units and force each unit to learn patterns that are more generalizable [25]. The dropout rate *p* serves as a tuning parameter that controls the intense of regularization. We applied dropout to all the hidden layers of D-GEX except for the outgoing edges from the input layer. The dropout rate was set to [0%, 10%, 25%] to compare the effect of different degrees of regularization.

2. *Momentum method* is a technique to accelerate gradient-based optimization. It accumulates a velocity in directions of gradients of the loss function across iterations and uses the velocity instead of the gradient to update parameters [12].

Given a loss function 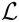 with respect to the parameters Θ of the neural network, the momentum is given by

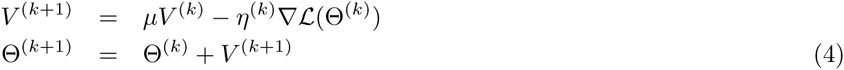

where *μ* ∈ [0,1] is the momentum coefficient, *η* is the learning rate, *V* is the velocity, and 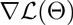 is the gradient of the loss function. Momentum method has been shown to improve the convergence rate particularly for training deep neural networks [12].

3. *Normalized initialization* is a technique to initialize the weights of deep neural networks [26]. The weights of a unit is sampled from a uniform distribution defined by,

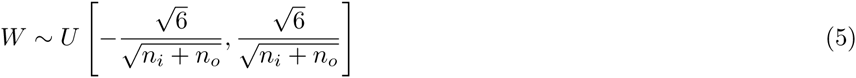

where *n_i_*, *n_o_* denote the number of fan-ins and fan-outs of the unit. It is designed to stabilize the variances of activation and back-propagated gradients during training [26]. The uniform distribution of the output layer of D-GEX was set to be within a smaller range of [−1 × 10^−4^, 1 × 10^−4^] as it was adopted with the linear activation function.

4. *Learning rate* was initialized to 5 × 10^−4^ or 3 × 10^−4^ depending on different architectures, and was decreased according to the training error on a subset of GEO-tr for monitoring the training process. Specifically, the training error was checked after each epoch, if the training error increased, the learning rate was multiplied by a decay factor of 0.9 until it reached a minimum learning rate of 1 × 10−^5^.

5. *Model selection* was performed based on GEO-va for the GEO data and 1000G-va for the GTEx data. Training was run for 200 epochs. The model was evaluated on GEO-va and 1000G-va after each epoch, and the model with the best performance was saved respectively.

D-GEX was implemented based on two Python libraries, Theano [27] and Pylearn2 [28]. Training was deployed on an Nvidia GTX TITAN Z graphics card with dual GPUs. The largest architecture of D-GEX (3 hidden layers with 9,000 hidden units in each hidden layer) contains ῀427 million parameters. Training half of the target genes with the largest architecture took around 6 hours. D-GEX is publicly available at https://github.com/uci-cbcl/D-GEX.

### 2.4 Linear regression

Linear regression (LR) for multi-task gene expression inference trains a model, 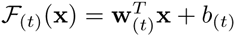, independently for each target gene *t*. 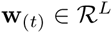, 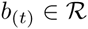 are the model parameters associated with each target gene *t*, and

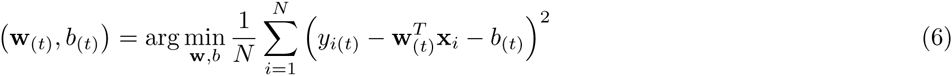

L1 or L2 penalties can be further introduced for regularization purpose. In these cases,

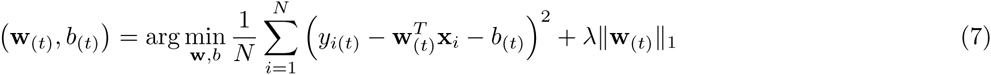

or

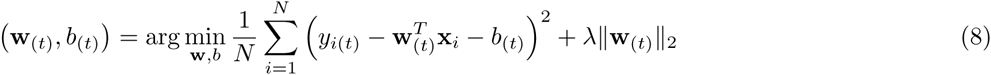

Linear regression (6) is currently adopted by the LINCS program. In our study, we evaluated both (6) and (7), (8) using scikit-learn [29]. The regularization parameter λ was tuned based on the performance on GEO-va and 1000G-va.

### 2.5 K-nearest neighbor regression

K-nearest neighbor (KNN) regression is a non-parametric and instance-based method. In standard KNN regression, a spatial data structure 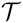 such as the KD tree [30] is built for training data in the feature space. Then, for any testing data, the k nearest training samples based on a certain distance metric are queried from 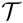. The average of their values is computed as the prediction.

However, the standard KNN regression may be biased when duplicated samples frequently exist in the data, such as the GEO microarray data. Therefore, in gene expression inference, a commonly adopted alternative is to query the k nearest genes rather than the k nearest samples. Specifically, for each target gene, its euclidean distances to all the landmark genes were calculated using the training samples. The k landmark genes with the least euclidean distances are determined as the k nearest landmark genes of the target gene. Then the average of their expression values in the testing samples is computed as the prediction for the target gene. Such algorithm is also consistent with the basic assumption of the LINCS program that, the expression of target genes can be computationally inferred from landmark genes. We call this algorithm the gene-based KNN (KNN-GE).

Due to the non-parametric and instance-based nature, KNN-GE does not impose any prior assumptions on the learning machine. Therefore, it is very flexible to model nonlinear patterns within the data. However, as performing prediction involves building and querying data structures that have to keep all the training data, KNN-GE suffers from poor scalability to growing data size and dimension. We evaluated KNN-GE in our study. The optimal k was selected based on the performance on GEO-va and 1000G-va.

## 3 Results

We have introduced two types of gene expression data, namely the GEO microarray data and the GTEx/1000G RNA-Seq data. We have formulated the gene expression inference as a multi-task regression problem, using the GEO data for training and both the GEO and the GTEx data for testing. We have also described our deep learning method D-GEX, and another two methods, linear regression and k-nearest neighbour regression, to solve the problem. Next, we show the predictive performances of the three methods on both the GEO data and the GTEx data.

### 3.1 Performance on the GEO data

D-GEX achieves the best performance on both GEO-va and GEO-te with 10% dropout rate (denoted as D-GEX-10%). Figure 1 and Table 1 show the overall performances of D-GEX-10% and the other methods on GEO-te. The complete performances of D-GEX with other dropout rates on both GEO-va and GEO-te are given in Supplementary Table S2 and S3. The largest architecture of D-GEX-10% (3 hidden layers with 9,000 hidden units in each hidden layer, denoted as D-GEX-10%-9000×3) achieves the best performance on both GEO-va and GEO-te. The relative improvements of D-GEX-10%-9000×3 are 15.33% over LR and 45.38% over KNN-GE. Besides D-GEX-10%-9000×3, D-GEX-10% consistently outperforms LR and KNN-GE on all the other architecture as shown in Figure 1. One possible explanation is that deep architectures enjoy much richer representability than shallow architectures, thus learning complex features is much easier from the perspective of optimization [31].

**Figure 1:**
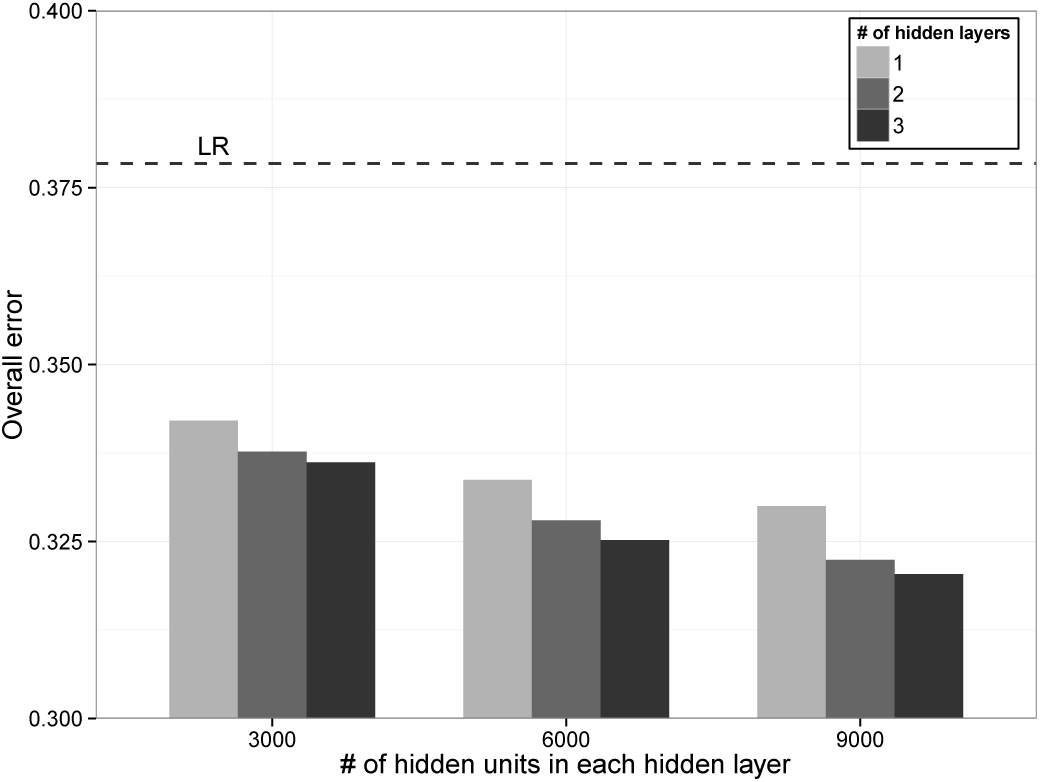
The overall errors of D-GEX-10% with different architectures on GEO-te. The performance of LR is also included for comparison.

**Table 1:**
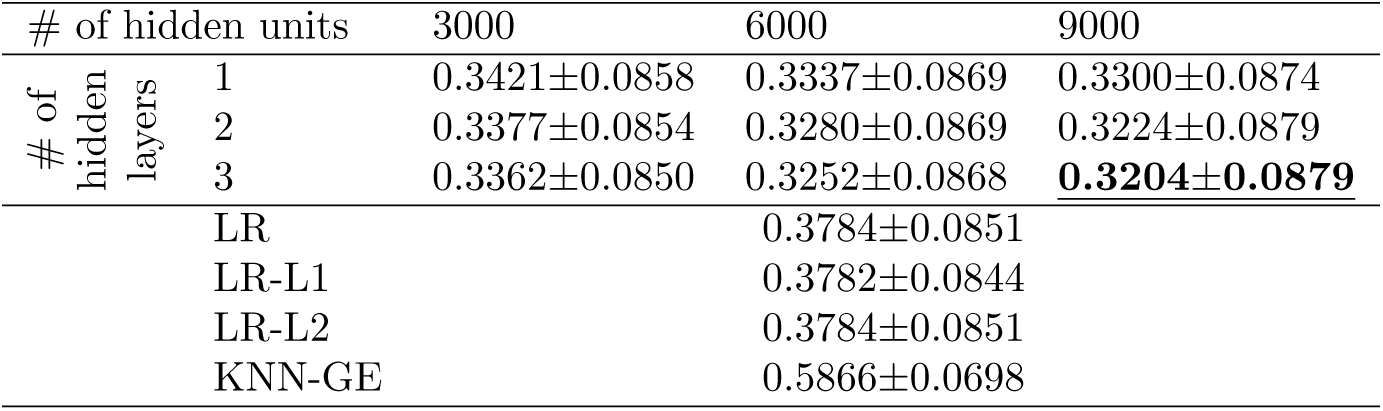
The overall errors of LR, LR-L1, LR-L2, KNN-GE and D-GEX-10% with different architectures on GEO-te. Numerics after “±” are the standard deviations of prediction errors over all target genes. The best performance of D-GEX-10% is shown in bold font. The performance selected using model selection by GEO-va of D-GEX-10% is underscored.

D-GEX also outperforms LR and KNN-GE for almost all of the target genes. Figure 2 shows the density plots of the predictive errors of all the target genes by LR, KNN-GE and GEX-10%-9000×3. Figure 3 shows a gene-wise comparative analysis between D-GEX-10%-9000×3 and the other two methods. D-GEX-10%-9000×3 outperforms LR in 99.97% of the target genes and outperforms KNN-GE in all the target genes. These results seem to suggest that D-GEX captured some intrinsic nonlinear features within the GEO data where LR and KNN-GE didn’t.

**Figure 2:**
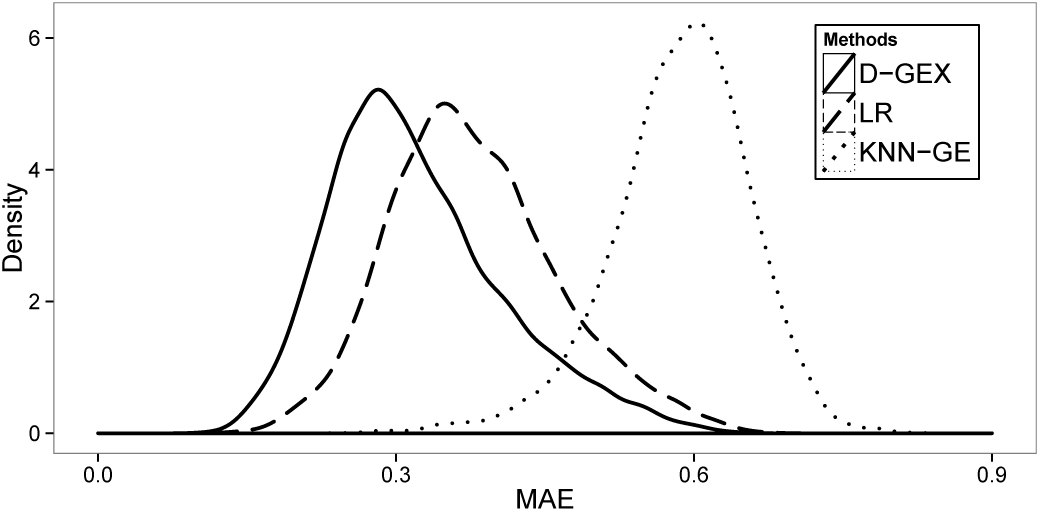
The density plots of the predictive errors of all the target genes by LR, KNN-GE and GEX-10%-9000×3 on GEO-te.

**Figure 3:**
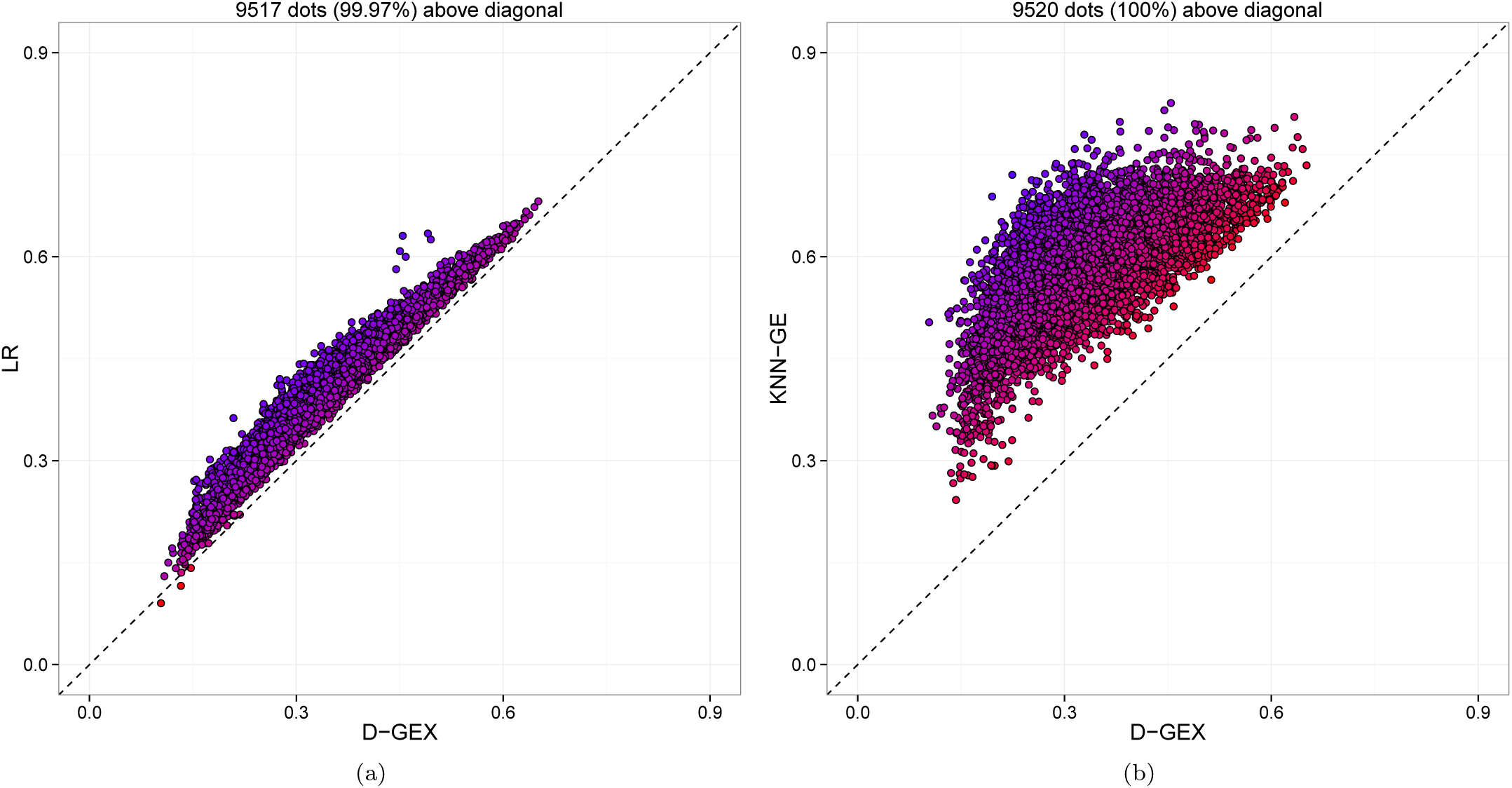
The predictive errors of each target gene by GEX-10%-9000×3 compared to LR and KNN-GE on GEO-te. Each dot represents 1 out of the 9,520 target genes. The x-axis is the MAE of each target gene by D-GEX, and the y-axis is the MAE of each target gene by the other method. Dots above diagonal means D-GEX achieves lower error compared to the other method. (a)D-GEX verse LR; (b)D-GEX verse KNN-GE.

Regularization methods do not improve LR significantly. Table 1 shows the relative improvements of LR-L1 and LR-L2 over LR are 0.05% and 0.00%. Thus, it is most likely that LR is underfitting which means linear model is not complex enough to represent the data. Therefore, regularization techniques that reduce model complexity are not helpful.

KNN-GE performs significantly worse than the other methods. One possible explanation is that the k nearest landmark genes for each target gene based on GEO-tr and GEO-te may not be fully consistent.

### 3.2 Performance on the GTEx data

Results on the GEO data demonstrate the significant improvement of D-GEX over LR and KNN-GE on the microarray platform. Yet in practice, the LINCS program trains regression models with the GEO data and performs gene expression inference on the L1000 data, which was generated with a different platform. Whether the significance of D-GEX preserves cross platforms requires further investigation. To explore the cross platforms scenario, we trained D-GEX with GEO-tr and evaluated its performances on GTEx-te which was generated with the RNA-Seq platform [20].

However, new challenges arise in this scenario as the intrinsic distributions of the training data and the testing data may be similar but not exactly equivalent. Particularly in gene expression profiling, discrepancies between microarray and RNA-Seq data have been systematically studied [32]. Such discrepancies bring specific challenges to deep learning as the complex features it learns in the training data may not generalize well to the testing data, which leads to overfitting and reduces the prediction power. Therefore, more aggressive regularization may be necessary for deep learning to retain the maximum commonality cross platforms while avoiding platform-dependent discrepancies.

D-GEX-25%-9000×2 (with 25% dropout rate, two hidden layers with 9000 hidden units in each layer) achieves the best performance on both 1000G-va and GTEx-te. The relative improvements of D-GEX-25%-9000×2 are 6.57% over LR and 32.62% over KNN-GE. Table 2 shows the overall performances of D-GEX-25% and the other methods on GTEx-te. The complete performances of D-GEX with other dropout rates on both 1000G-va and GTEx-te are given in Supplementary Table S4 and S5.

**Table 2:** The overall errors of LR, LR-L1, LR-L2, KNN-GE and D-GEX-25% with different architectures on GTEx-te. Numerics after “±” are the standard deviations of prediction errors over all target genes. The best performance of D-GEX-25% is shown in bold font. The performance selected using model selection by 1000G-va of D-GEX-25% is underscored.

| # of hidden units |  | 3000 | 6000 | 9000 |
| --- | --- | --- | --- | --- |
| # of hidden layers | 1 | 0.4507±0.1231 | 0.4428±0.1246 | 0.4394±0.1253 |
| | 2 | 0.4586±0.1194 | 0.4446±0.1226 | **<u>0.4393±0.1239</u>** |
| | 3 | 0.5160±0.1157 | 0.4595±0.1186 | 0.4492±0.1211 |
| | LR | | 0.4702±0.1234 | |
| | LR-L1 | | 0.5667±0.1271 | |
| | LR-L2 | | 0.4702±0.1234 | |
| | KNN-GE | | 0.6520±0.0982 | |

D-GEX still outperforms LR and KNN-GE in most of the target genes. Figure 4 also shows the gene-wise comparative analysis between D-GEX-25%-9000×2 and the other two methods. D-GEX-25%-9000×2 outperforms LR in 81.31% of the target genes and outperforms KNN-GE in 95.54% of the target genes. Therefore, the significance of D-GEX on the microarray platform basically preserves on the RNA-Seq platform. However, unlike the results on the GEO data, there is a noticeable number of target genes that D-GEX gets higher error than the other methods on the GTEx data. Thus, the expression patterns of these target genes D-GEX learned on the GEO data may be platform dependent and do not generalize well to the GTEx data. It is noteworthy that although the general performance of KNN-GE is still poor on the GTEx data, its errors on some of the target genes are significantly lower than D-GEX (dots in bottom right part of Figure 4(b)). This is likely due to the gene-based aspect of KNN-GE that the numerical values predicted on target genes were not computed based on GEO-tr but based on GTEx-te itself. Therefore, the expression patterns captured by KNN-GE may be cross platforms invariant.

**Figure 4:**
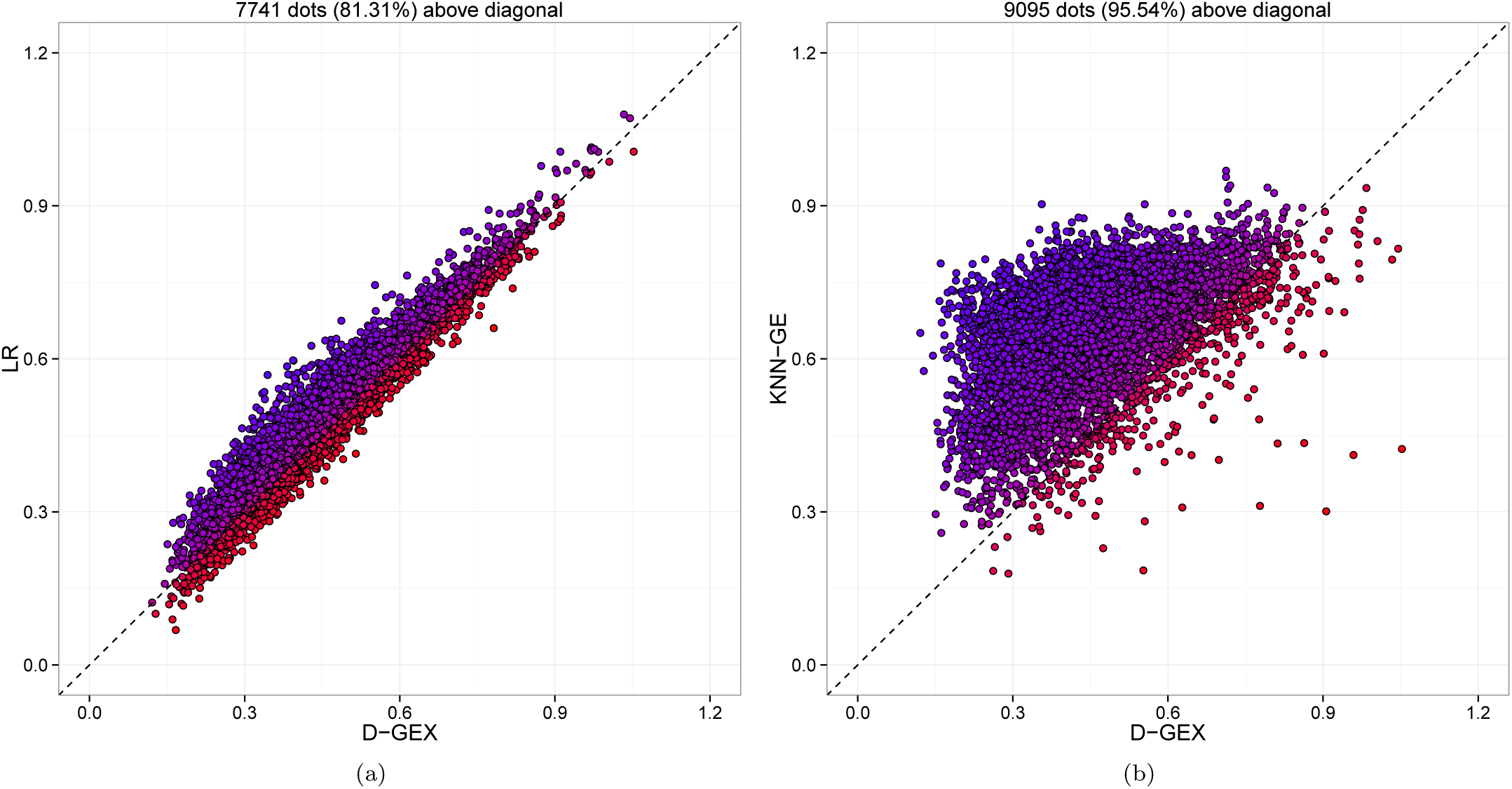
The predictive errors of each target gene by GEX-25%-9000×2 compared to LR and KNN-GE on GTEx-te. Each dot represents 1 out of the 9,520 target genes. The x-axis is the MAE of each target gene by D-GEX, and the y-axis is the MAE of each target gene by the other method. Dots above diagonal means D-GEX achieves lower error compared to the other method. (a)D-GEX verse LR; (b)D-GEX verse KNN-GE.

Dropout regularization effectively improves the performance of D-GEX on the GTEx data as shown in Figure 5. Without dropout, the overall error of D-GEX-9000×2 on GTEx-te slightly decreases at the beginning of training and then quickly increases, clearly implying overfitting. However, with 25% dropout rate, D-GEX-9000×2 achieves the best performance on both 1000G-va and GTEx-te.

**Figure 5:**
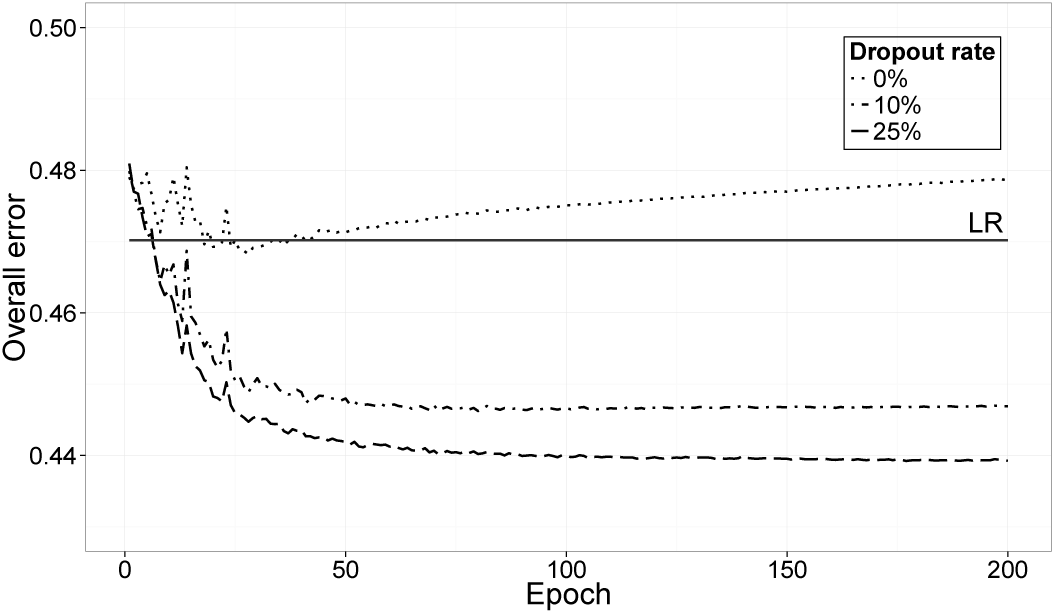
The overall error decreasing curves of D-GEX-9000×2 on GTEx-te with differant dropout rates. The x-axis is the training epoch and the y-axis is the overall error. The overall error of LR is also included for comparison.

### 3.3 Interpreting the learned neural network

We have demonstrated the performance of our deep learning method D-GEX on both the GEO microarray data and the GTEx RNA-Seq data. D-GEX outperforms linear regression on both types of expression data. On the other hand, interpreting the learned linear model from linear regression is straightforward as coefficients with large absolute value indicate strong dependencies between landmark genes and target genes. But for deep learning, currently there are no established methods to interpret the neutral networks learned from gene expression data. Next, we attempt to explore the learned neural networks with two strategies, a) visualizing the major weights of the learned neural networks and b) examining the nonlinearity captured by the hidden layers.

1. *Visualizing the major weights* is a strategy inspired by the method of interpreting linear model that coefficients with large absolute value indicate strong dependencies between inputs and targets. Similarly, we examined the weights of the learned neural network of D-GEX-10%-3000x 1 that was trained based on half of the target genes of GEO-tr and GEO-va. The weights from input to hidden units were randomly initialized with dense connections. However, after learning, the connections became so sparse that each input unit was primarily connected to only a few hidden units with the weights to the rest of hidden units decayed to near zero. Similar patterns were also observed for connections from the hidden to the output layer. Therefore, we created a visualization map of the learned connections by removing those with weights near zeros. Specifically, for each input unit (landmark gene), we calculated the mean and the standard deviation of the weights of the connections between the input unit and the 3,000 hidden units. Then we only retained the major weights that were 4 standard deviations away from the mean. Likewise, we used a threshold of 5 standard deviations to retain the major weights of the connections between the output units (target genes) and the hidden units. We colored the weights differently so that red indicates positive weights and blue indicates negative weights. Supplementary Figure S3 shows the final visualization map. From the visualization map, we noticed two interesting observations: a) Most of the units in the input layer and the output layer have connections to the hidden layer. In contrast, only a sparse number of units in the hidden layer have connections to the input and the output layer. Specially, the connections to the output layer are dominated by a few hidden units, which we refer to as the “hub units”. b) Lots of the “hub units” seem to have only one type of connections to the output layer, e.g. some of them only have positive connections (red edges), while some other units only have negative connections (blue edges). It seems that these “hub units” may have captured some strong local correlations between the landmark genes and target genes.

2. *Examining the nonlinearity* is a strategy to show that the intermediate hidden layers have captured some nonlinearity within the raw expression data. The neural networks we used are quite complex, containing several layers and many hidden units, each of which is activated through a nonlinear transfer function. To dissect the nonlinear contribution, we took a relatively simple approach by focusing on the representation (activations) from the last hidden layer. Each of the hidden unit in that layer can be viewed as a feature generated through some nonlinear transformation of the landmark genes. We then studied whether a linear regression based on these nonlinear features can achieve better performance than a linear regression based solely on the landmark genes. For this purpose, we measured the linear correlation between the activations from the last hidden layer of D-GEX-10%-9000×3 and the final targets (the expression of target genes), and compared it with the linear correlation between the raw inputs and the final targets. Normally, coefficient of determination (*R*^2^) is used to compare the fitnesses of different linear models. Since the dimensionality has changed from the raw inputs to the transformed activations, we used adjusted *R*^2^ [33] to specifically account for the change in dimensionality. We calculated the adjusted *R*^2^ of both the raw inputs and the transformed activations for each target gene based on GEO-tr. Supplementary Figure S2 shows the gene-wise comparison of adjusted *R*^2^ between the raw inputs and the transformed activations. The transformed activations have a larger adjusted *R*^2^ than the raw inputs in 99.99% of the target genes. It seems to indicate that the intermediate hidden layers have systematically captured some nonlinearity within the raw expression data that would be ignored by simple linear regression. After the nonlinear transformation through the hidden layers, the activations fit the final targets significantly better than the raw inputs using a simple linear model. The analysis seems to suggest that most of the target genes benefit from the additional nonlinear features, although to a different extent as characterized by the adjusted *R*^2^.

### 3.4 Inference on the L1000 data

The LINCS program has used the L1000 technology to measure the expression profiles of the 978 landmark genes under a variety of experimental conditions. It currently adopts linear regression to infer the expression values of the 21,290 target genes based on the GEO data. We have demonstrated our deep learning method D-GEX achieved significantly improvement on prediction accuracy over linear regression on the GEO data. Therefore, we have re-trained GEX-10%-9000×3 using all the 978 landmark genes and the 21,290 target genes from the GEO data and inferred the expression values of unmeasured target genes from the L1000 data. The full dataset consists of 1,328,098 expression profiles and can be downloaded at https://cbcl.ics.uci.edu/public_data/D-GEX/l1000_n1328098x22268.gctx. We hope this dataset will be of great interest to researchers who are currently querying the LINCS L1000 data.

## 4 Discussion

Revealing the complex patterns of gene expression under numerous biological states requires both cost-effective profiling tools and powerful inference frameworks. While the L1000 platform adopted by the LINCS program can efficiently profile the ῀1,000 landmark genes, the linear-regression-based inference does not fully leverage the nonlinear features within gene expression profiles to infer the ῀21,000 target genes. We presented a deep learning method for gene expression inference that significantly outperforms linear regression on the GEO microarray data. With dropout as regularization, our deep learning method also preserves cross platforms generalizability on the GTEx RNA-Seq data. In summary, deep learning provides a better model than linear regression for gene expression inference. We believe it achieves more accurate predictions for target gene expressions of the LINCS dataset generated from the L1000 platform.

Interpreting the internal representation of deep architectures is notoriously difficult. Unlike other machine learning tasks such as computer vision, where we can visualize the learned weights of hidden units as meaningful image patches, interpreting the deep architectures learned by biological data requires novel thinking. We attempted to interpret the internal structures of the neural networks learned from gene expression data using strategies that were inspired by linear model. Yet, more systematic studies may require advanced computational frameworks that are specifically designed for deep learning. Unsupervised feature learning methods, such as autoencoder [34] and restricted Boltzmann machine [35] may provide some insights on this problem.

In the current setting, target genes were randomly partitioned into multiple sets, and each set was trained separately using different GPUs due to hardware limitations. Alternatively, we could first cluster target genes based on their expression profiles, and then partition them accordingly rather than randomly. The rationale is that target genes sharing similar expression profiles share weights in the context of multi-task neural networks. Ultimately, the solution is to jointly train all target genes, either by using GPUs with larger memory such as the more recent Nvidia Tesla K80, or by exploiting multi-GPU techniques [9].

## References

[1] Justin Lamb, Emily D Crawford, David Peck, Joshua W Modell, Irene C Blat, Matthew J Wrobel, Jim Lerner, Jean-Philippe Brunet, Aravind Subramanian, Kenneth N Ross, et al. The connectivity map: using gene-expression signatures to connect small molecules, genes, and disease. science, 313(5795):1929–1935, 2006.

[2] Mukesh Bansal, Vincenzo Belcastro, Alberto Ambesi-Impiombato, and Diego Di Bernardo. How to infer gene networks from expression profiles. Molecular systems biology, 3(1), 2007.

[3] David Peck, Emily D Crawford, Kenneth N Ross, Kimberly Stegmaier, Todd R Golub, and Justin Lamb. A method for high-throughput gene expression signature analysis. Genome biology, 7(7):R61, 2006.

[4] Ron Edgar, Michael Domrachev, and Alex E Lash. Gene expression omnibus: Ncbi gene expression and hybridization array data repository. Nucleic acids research, 30(1):207–210, 2002.

[5] Jeff Hasty, David McMillen, Farren Isaacs, and James J Collins. Computational studies of gene regulatory networks: in numero molecular biology. Nature Reviews Genetics, 2(4):268–279, 2001.

[6] Guibo Ye, Mengfan Tang, Jian-Feng Cai, Qing Nie, and Xiaohui Xie. Low-rank regularization for learning gene expression programs. PloS one, 8(12):e82146, 2013.

[7] Yoshua Bengio, Aaron Courville, and Pascal Vincent. Representation learning: A review and new perspectives. Pattern Analysis and Machine Intelligence, IEEE Transactions on, 35(8):1798–1828, 2013.

[8] Dan Ciresan, Ueli Meier, and Jürgen Schmidhuber. Multi-column deep neural networks for image classification. In Computer Vision and Pattern Recognition (CVPR), 2012 IEEE Conference on, pages 3642–3649. IEEE, 2012.

[9] Adam Coates, Brody Huval, Tao Wang, David Wu, Bryan Catanzaro, and Ng Andrew. Deep learning with cots hpc systems. In Proceedings of The 30th International Conference on Machine Learning, pages 1337–1345, 2013.

[10] Geoffrey E Hinton, Nitish Srivastava, Alex Krizhevsky, Ilya Sutskever, and Ruslan R Salakhutdinov. Improving neural networks by preventing co-adaptation of feature detectors. arXiv preprint arXiv:1207.0580, 2012.

[11] Pierre Baldi and Peter J Sadowski. Understanding dropout. In Advances in Neural Information Processing Systems, pages 2814–2822, 2013.

[12] Ilya Sutskever, James Martens, George Dahl, and Geoffrey Hinton. On the importance of initialization and momentum in deep learning. In Proceedings of the 30th International Conference on Machine Learning (ICML-13), pages 1139–1147, 2013.

[13] Alex Krizhevsky, Ilya Sutskever, and Geoffrey E Hinton. Imagenet classification with deep convolutional neural networks. In Advances in neural information processing systems, pages 1097–1105, 2012.

[14] Richard Socher, Cliff C Lin, Chris Manning, and Andrew Y Ng. Parsing natural scenes and natural language with recursive neural networks. In Proceedings of the 28th international conference on machine learning (ICML-11), pages 129–136, 2011.

[15] Geoffrey Hinton, Li Deng, Dong Yu, George E Dahl, Abdel-rahman Mohamed, Navdeep Jaitly, Andrew Senior, Vincent Vanhoucke, Patrick Nguyen, Tara N Sainath, et al. Deep neural networks for acoustic modeling in speech recognition: The shared views of four research groups. Signal Processing Magazine, IEEE, 29(6):82–97, 2012.

[16] P Baldi, P Sadowski, and D Whiteson. Searching for exotic particles in high-energy physics with deep learning. Nature communications, 5, 2014.

[17] Pietro Di Lena, Ken Nagata, and Pierre Baldi. Deep architectures for protein contact map prediction. Bioinformatics, 28(19):2449–2457, 2012.

[18] Michael KK Leung, Hui Yuan Xiong, Leo J Lee, and Brendan J Frey. Deep learning of the tissue-regulated splicing code. Bioinformatics, 30(12):i121–i129, 2014.

[19] Daniel Quang, Yifei Chen, and Xiaohui Xie. Dann: a deep learning approach for annotating the pathogenicity of genetic variants. Bioinformatics, page btu703, 2014.

[20] John Lonsdale, Jeffrey Thomas, Mike Salvatore, Rebecca Phillips, Edmund Lo, Saboor Shad, Richard Hasz, Gary Walters, Fernando Garcia, Nancy Young, et al. The genotype-tissue expression (gtex) project. Nature genetics, 45(6):580–585, 2013.

[21] Kristin G Ardlie, David S Deluca, Ayellet V Segrè, Timothy J Sullivan, Taylor R Young, Ellen T Gelfand, Casandra A Trowbridge, Julian B Maller, Taru Tukiainen, Monkol Lek, et al. The genotype-tissue expression (gtex) pilot analysis: Multitissue gene regulation in humans. Science, 348(6235):648–660, 2015.

[22] Tuuli Lappalainen, Michael Sammeth, Marc R Friedländer, Peter ACt Hoen, Jean Monlong, Manuel A Rivas, Mar Gonzàlez-Porta, Natalja Kurbatova, Thasso Griebel, Pedro G Ferreira, et al. Transcriptome and genome sequencing uncovers functional variation in humans. Nature, 501(7468):506–511, 2013.

[23] David E Rumelhart, Geoffrey E Hinton, and Ronald J Williams. Learning representations by back-propagating errors. Cognitive modeling, 5, 1988.

[24] Yifei Chen. Machine Learning for Large-Scale Genomics: Algorithms, Models and Applications. PhD thesis, University of California, Irvine, ProQuest, UMI Dissertations Publishing, 12 2014.

[25] Nitish Srivastava, Geoffrey Hinton, Alex Krizhevsky, Ilya Sutskever, and Ruslan Salakhutdinov. Dropout: A simple way to prevent neural networks from overfitting. The Journal of Machine Learning Research, 15(1):1929–1958, 2014.

[26] Xavier Glorot and Yoshua Bengio. Understanding the difficulty of training deep feedforward neural networks. In International conference on artificial intelligence and statistics, pages 249–256, 2010.

[27] James Bergstra, Olivier Breuleux, Frédéric Bastien, Pascal Lamblin, Razvan Pascanu, Guillaume Desjardins, Joseph Turian, David Warde-Farley, and Yoshua Bengio. Theano: a CPU and GPU math expression compiler. In Proceedings of the Python for Scientific Computing Conference (SciPy), June 2010.

[28] Ian J Goodfellow, David Warde-Farley, Pascal Lamblin, Vincent Dumoulin, Mehdi Mirza, Razvan Pascanu, James Bergstra, Frédéric Bastien, and Yoshua Bengio. Pylearn2: a machine learning research library. arXiv preprint arXiv:1308.4214, 2013.

[29] F. Pedregosa, G. Varoquaux, A. Gramfort, V. Michel, B. Thirion, O. Grisel, M. Blondel, P. Prettenhofer, R. Weiss, V. Dubourg, J. Vanderplas, A. Passos, D. Cournapeau, M. Brucher, M. Perrot, and E. Duchesnay. Scikit-learn: Machine learning in Python. Journal of Machine Learning Research, 12:2825–2830, 2011.

[30] Jon Louis Bentley. Multidimensional binary search trees used for associative searching. Communications of the ACM, 18(9):509–517, 1975.

[31] Yoshua Bengio. Learning deep architectures for AI. Foundations and trends^®^ in Machine Learning, 2(1):1–127, 2009.

[32] Charles Wang, Binsheng Gong, Pierre R Bushel, Jean Thierry-Mieg, Danielle Thierry-Mieg, Joshua Xu, Hong Fang, Huixiao Hong, Jie Shen, Zhenqiang Su, et al. The concordance between rna-seq and microarray data depends on chemical treatment and transcript abundance. Nature biotechnology, 32(9):926–932, 2014.

[33] Henry Theil. Economic forecasts and policy. 1958.

[34] Pascal Vincent, Hugo Larochelle, Yoshua Bengio, and Pierre-Antoine Manzagol. Extracting and composing robust features with denoising autoencoders. In Proceedings of the 25th international conference on Machine learning, pages 1096–1103. ACM, 2008.

[35] Geoffrey Hinton. A practical guide to training restricted boltzmann machines. Momentum, 9(1):926, 2010.

